# Temperature-dependent changes to host-parasite interactions alter the thermal performance of a bacterial host

**DOI:** 10.1101/554717

**Authors:** Daniel Padfield, Meaghan Castledine, Angus Buckling

## Abstract

Thermal performance curves (TPCs) are used to predict changes in species interactions, and hence range shifts, disease dynamics and community composition, under forecasted climate change. Species interactions might in turn affect TPCs. Here, we investigate whether temperature-dependent changes in a microbial host-parasite interaction (the bacterium *Pseudomonas fluorescens*, and its bacteriophage, SBWΦ2) changes the host TPC. The bacteriophage had a narrower infectivity range, with their critical thermal maximum ∼6°C lower than those at which the bacteria still had high growth. Consequently, in the presence of phage, the host TPC had a higher optimum temperature and a lower maximum growth rate. These changes were driven by a temperature-dependent evolution, and cost, of resistance; the largest cost of resistance occurring where bacteria grew best in the absence of phage. Our work highlights how ecological and evolutionary mechanisms can alter the effect of a parasite on host thermal performance, even over very short timescales.

**Data accessibility statement:** All data and R code used in the analysis will be made available on GitHub and archived on Zenodo.

## Introduction

An often overlooked concern surrounding climate change is its impacts on host-parasite interactions^1^. The effect of temperature on such interactions is likely widespread, as temperature influences the physiology, ecology and evolution of both hosts and parasites^2–5^. However, the sign and strength of the effects of warming on host-parasite interactions will be context dependent, changing with the host, parasite and environmental conditions in question. One approach to predict the potential impacts of warming on host-parasite interactions has been based around thermal performance curves (TPCs) of, and differences between, key host and parasite traits^2,6,7^. For example, it has been argued that as hosts generally have a narrower thermal range and lower thermotolerance than their parasites^8–10^, they are more susceptible to disease at temperatures further away from their optimum temperature.

An inevitable consequence of temperature dependent changes in host-pathogen interactions^11^ is a change in the host’s TPC in the presence, versus the absence, of the parasite. For example, if the largest impact of the parasite occurs at the host’s optimum growth temperature, key traits such as maximum growth rate and optimum temperature of the host could change. In addition to the ecological feedbacks resulting from differences in the thermal performance of host and parasite traits, rapid (co)evolution of resistance and infectivity traits could play a major role in altering TPCs. Crucially, TPCs of hosts and parasites are typically assumed to be fixed across time and in different abiotic and biotic environments^7,8,12,13^, but the presence of a predator can alter the TPC of the prey^14^ and the prey’s evolutionary response to warming^15^. If parasites affect the thermal performance of their host, this may alter some of the predictions of range shifts and disease dynamics expected under climate change.

To date, most experimental and theoretical work on the thermal performance of organisms is done on single species, where naturally occurring parasites, symbionts and microbiota are greatly or completely removed^16–18^. Consequently, it is unknown if parasites alter the TPC of host fitness and influence key traits such as the optimal, *T*_*opt*_, and cardinal (critical thermal maximum, *CT*_*max*_, and minimum, *CT*_*min*_) temperatures of host growth. Understanding these potential impacts is critical to assess the effect of climate change on ecological and evolutionary dynamics of host-parasite pairs, as well as predicting the consequences of novel host-parasite interactions that will occur in a warmer world. Here, we explicitly determine how and why interactions with a parasite affect host thermal performance in arguably the most common host-pathogen interaction on the planet: that between bacteria and their viruses (bacteriophage).

We focus on a well-studied system, the bacterium *Pseudomonas fluorescens* SBW25 and its phage, *SBW*Ф2. This system has been used extensively for studying bacteria-virus ecological and evolutionary interactions^19–22^. Over a wide range of temperatures, we measured the replication rate of the phage and the growth rate of the bacteria in the presence and absence of the phage. We utilised the ‘traits’ that underpin TPCs to compare biologically meaningful parameters. We hypothesised that any large difference in thermal performance of bacteria and phage would change the thermal performance of bacteria in the presence *vs.* the absence of phage. Given the importance of evolution occurring over ecological timescales, especially in microbial populations with large population sizes and short generation times, we also investigated evolutionary changes in host populations to determine whether resistance evolution explained any changes in host thermal performance.

## Results

### Bacteria and phage had mismatches in their thermal performance

We measured phage replication rate and bacterial growth rate across eight temperature (15 – 37 °C) to determine whether there were mismatches in the thermal performance of the host and its parasite. To do this, we modelled the thermal performance curve of each rate and used estimated and derived parameters of the model (see Equation 2 in Methods) as traits that we used to compare the thermal responses of bacteria and phage. Phage replication rate increased to a thermal optimum, *T*_*opt*_, of 27 °C (95% credible intervals [CI]: 26.5 – 27.5 °C) before rapidly declining to a negative replication rate by 30 °C (Fig. 1a). The critical thermal maximum, *CT*_*max*_, of phage replication was 29.2 °C (95% CI: 29.0 – 29.4 °C), beyond which phage decreased in abundance over 4 hours (Fig. 1a). This indicated that phage struggled to infect the host at temperatures beyond their *T*_*opt*_, which was similar to previous work that measured the coevolution of this bacteria-phage system across temperatures^22^. The bacteria, *Pseudomonas fluorescens*, had a similar optimum temperature (Fig. 1b [blue]; *T*_*opt*_ = 28 °C; 95% CI: 27.1 – 29.0 °C), but growth was maintained well beyond *T*_*opt*_, with high growth rates still occurring at 35 °C (Fig. 1b), > 6 °C above the *CT*_*max*_ of the phage. This could act as a high temperature refuge for the bacteria as phage infection at these temperatures is extremely low. Due to these thermal mismatches between the infectivity range of phage and the growth range of bacteria, it was expected that the parasite would alter the thermal performance of its host.

**Figure 1.**
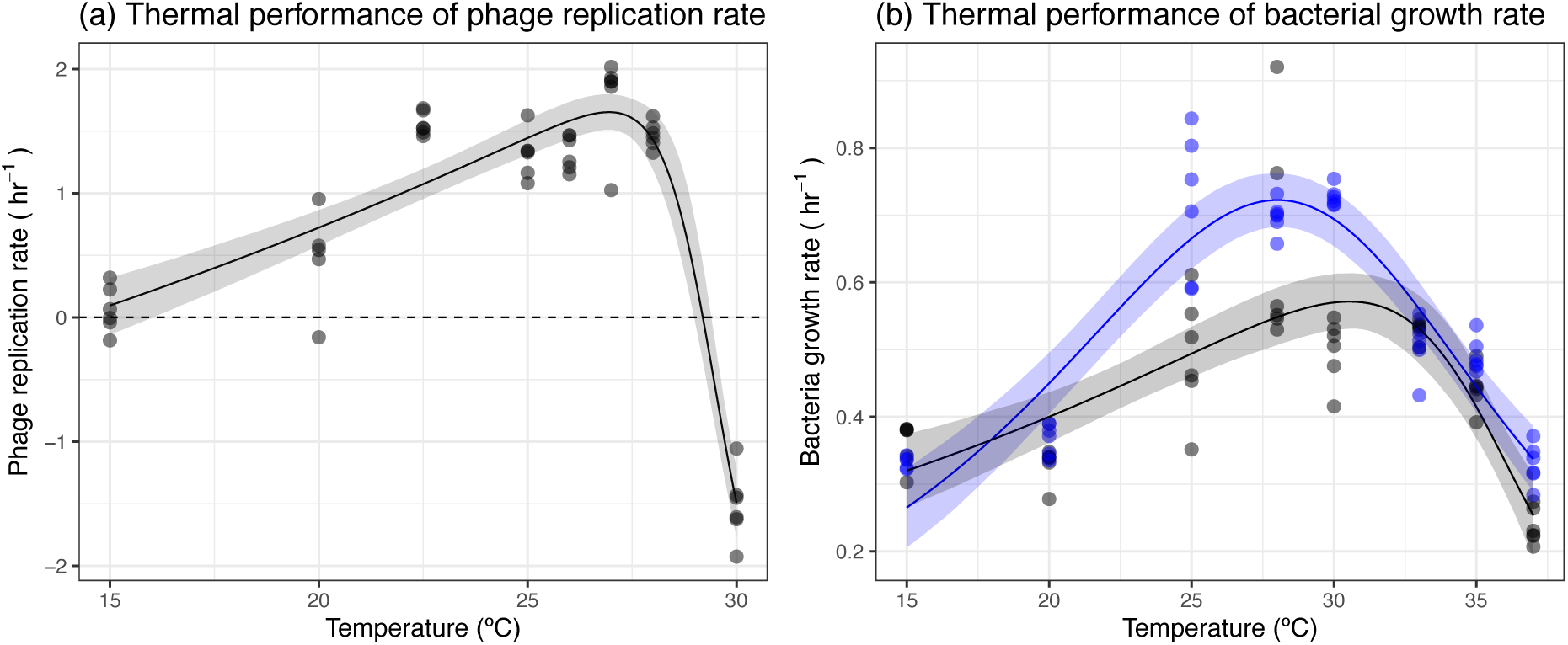
Thermal performance of phage and bacteria. (a) Phage replication increases with temperature up to an optimum of before declining rapidly to a negative replication rate at 30 °C. (b) Bacteria growth shows unimodal responses to temperature in the presence (black) and absence of phage (blue). However, phage changed the shape of the thermal response. Points represent an independent replicate at each temperature. Solid lines represent the mean prediction and shaded bands represent the 95% credible interval of predictions. In (a) the dashed line represents 0 growth, below which phage abundance decreased.

### Phage altered the thermal performance of its bacterial host

Due to the thermal mismatches between bacteria and phage, we explored whether phage altered the thermal performance of its host. To do this, we measured the growth rate of bacteria in the presence and absence of the phage and compared key traits that underpinned the thermal performance curve (see Methods). We observed marked differences in the response of bacteria to temperature when in the presence of its phage (Fig. 1b & Table S1). Phage increased the optimum temperature of bacterial growth (Fig. 2c), shifting *T*_*opt*_ from 28 °C (95% CI: 27.1 – 29.0 °C) to 30.6 °C (95% CI: 29.0 – 32.1 °C). This shift in *T*_*opt*_ resulted in a 20.1% (95% CI: 13 - 27.3%) decline in the maximal growth rate, *r*_*max*_, in the presence of phage (Fig. 3d), as the largest impacts of phage on bacterial growth occurred at temperatures where growth was highest (Fig. 1b & Fig. 2a). To better understand the non-linear, temperature dependent effect of phage on bacterial growth, we calculated the relative fitness of bacteria in the presence of phage across temperatures (see Methods; Fig. 2a). Differences were greatest at intermediate temperatures where growth in the absence of phage was fastest (Fig. 2a, where relative fitness was <1), whereas no significant change in growth rate was observed at the low and high temperatures measured (credible intervals of predictions overlap 1). The non-linear changes to bacterial growth also resulted in differences in other key traits (Table S1) such as the activation energy (*E*; Fig. 3b) and the deactivation energy (*E*_*h*_; Fig. 3e).

**Figure 2.**
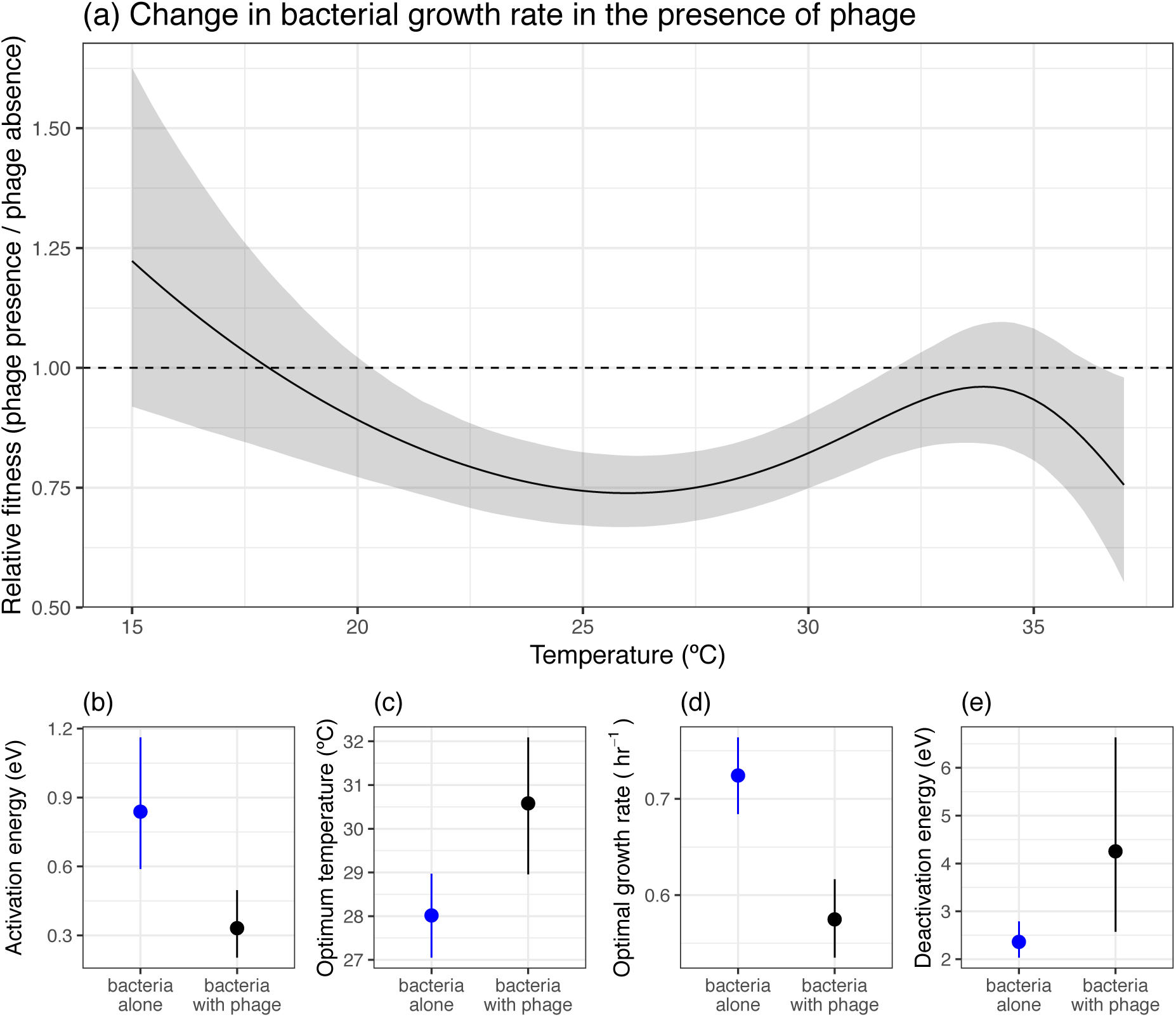
Effect of phage on the thermal performance of bacteria. (a) Phage altered the growth rate of bacteria (calculated as relative fitness) in a non-linear fashion with increasing temperatures. (b-e) The effect of phage on key thermal performance traits. Phage altered the (b) activation energy, (c) optimum temperature, (d) optimal growth rate and (e) deactivation energy. In (a) the solid line represents the mean prediction and shaded band represents the 95% credible interval of predictions. The dashed line at y = 1 would indicate that phage do not alter growth rate. Below 1, phage reduces the growth rate of the bacteria. In (b-e) points and lines represent the mean and 95% credible intervals of the estimated parameters.

**Figure 3.**
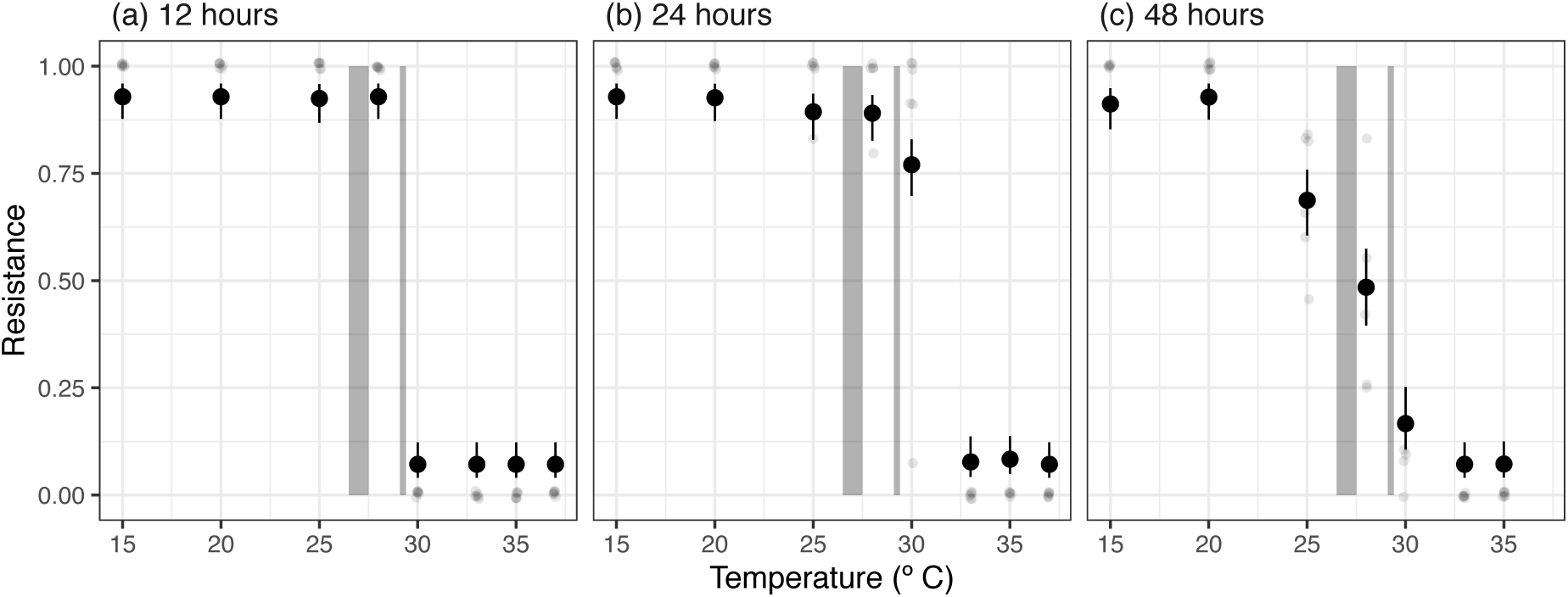
Levels of resistance of *Pseudomonas fluorescens* to phage through time and across temperatures. After 12 hours, populations are completely resistant at temperatures of 28 °C or lower. After 24 hours, most bacteria populations at 30 °C, close to the estimated critical thermal maxima of the phage, have evolved resistance, but populations beyond the infectivity range of the phage remain susceptible. After 48 hours, at temperatures where phage impact bacterial growth, intermediate levels of resistance are observed. Small points represent the observed level of resistance for a population. Large points represent the predicted levels of resistance from a binomial regression with 95% confidence intervals. Shaded regions represent the upper and lower credible intervals of the optimum temperature and critical thermal maxima of the phage.

### The evolution and cost of resistance was temperature dependent

It is possible that the change in thermal performance of *Pseudomonas fluorescens* could have resulted simply from the mismatches in thermal performances of the host and parasite. Up to *T*_*opt*_ of the phage (∼27 °C), phage infection reduced the abundance and thus population growth rate of the bacteria. Consequently, the rapid decline of phage replication at temperatures above 30 °C, while bacteria still had high growth rates, could explain the shift to a higher *T*_*opt*_ of the bacteria in the presence of phage. However, bacteria can rapidly evolve resistance to phage within the timescales of our assays, and this has been demonstrated in our host-parasite pair^23,24^. If, as expected, resistance is costly, and resistance does not evolve at temperatures beyond the phage *CT*_*max*_, the effect of phage on the thermal performance of the host may in part be driven by evolutionary change. To investigate this, we measured the resistance of bacteria through a single logistic growth curve at each temperature (Fig. 3). The evolution of phage resistance changed across temperatures and through time, and there was a significant time x temperature interaction (likelihood ratio test comparing models with and without time x temperature interaction: Δ*d.f.* = 13, *F* = 11.56, *P* < 0.001). There was no measurable resistance in the ancestral bacteria, but after just 12 hours, all populations at 28 °C (close to *T*_*opt*_ of phage replication [∼27 °C]) or lower were close to 100% resistant (Fig. 3a), consistent with a selective sweep. In contrast, resistance rarely, or never, evolved at temperatures well above those of the critical thermal maximum of phage replication rate (33 °C and higher, Fig. 4). Where resistance did evolve at these temperatures, it was at very low frequency (1 clone out of 12). We found no bacteria still living at 37 °C after 48 hours, indicating that although growth occurs at those temperatures, this is quickly proceeded by death.

**Figure 4.**
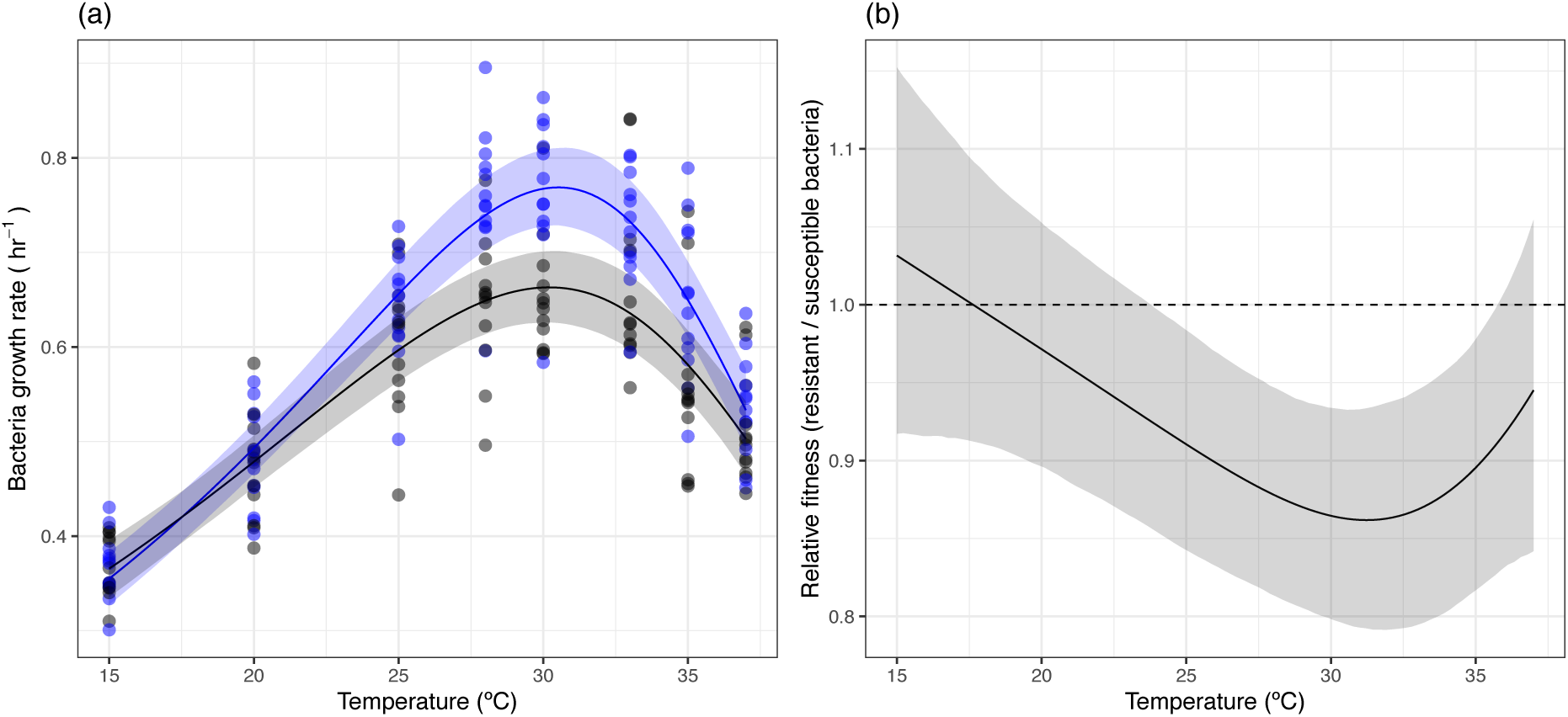
Temperature dependent cost of resistance in *Pseudomonas fluorescens*. (a) The thermal performance of susceptible (blue) and resistant (black) clones. Resistant clones have a lower maximum growth rate. (b) The derived selection coefficient of resistance across temperatures. The cost of resistance changes across temperatures, being greatest at 30 °C and other temperatures where growth in the absence of phage is high. In (a) points represent individual clones, solid lines represent the mean prediction and shaded bands represent the 95% credible interval of predictions. In (a) the solid line represents the mean prediction and shaded band represents the 95% credible interval of predictions. The dashed line at y = 1 would indicate that phage do not alter growth rate. Below 1, phage reduces the growth rate of the bacteria.

At temperatures where phage altered the growth rate of bacteria (25, 28 & 30 °C), we observed significant changes in the proportion of resistance through time (see Table S2 for pairwise differences of resistance through time for each temperature). Resistance evolved and was at high proportions after 12 or 24 hours where populations were still in exponential growth phase. However, after 48 hours, when populations had reached stationary phase at all temperatures apart from 15 and 20 °C (Figure S3), the proportion of resistance decreased significantly (Fig. 4c). From 24 to 48 hours, 25 °C resistance fell from 0.89 (95% CI: 0.83 - 0.94) to 0.69 (95% CI: 0.60 – 0.78), at 28 °C from 0.89 (95% CI: 0.83 - 0.93) to 0.48 (95% CI: 0.40 - 0.57) and at 30 °C from 0.77 (95% CI 0.68 – 0.83) to 0.17 (95% CI: 0.11 - 0.25). This temporal effect did not occur at low and high temperatures where there was little effect of phage on bacterial growth rate (Fig. 2a & Table S2), suggesting that there was a non-linear cost of resistance across the temperature range.

To confirm whether there was a cost of resistance and if any cost varied with temperature, we isolated three clones from 12 independent replicates that were grown for 12 hours at 28 °C with and without phage to produce resistant and susceptible clones. After checking for resistance, we measured the thermal performance of resistant and susceptible clones in the absence of phage. The thermal performance of resistant clones differed from that of susceptible clones (Fig. 4), closely matching the patterns observed when bacteria were grown with phage (Fig. 1b & Fig. 2a). At low and high temperatures, there were no differences in the growth rate of resistant and susceptible clones (Fig. 4). However, at temperatures where growth of susceptible clones was highest (25 – 30 °C), there was a cost of resistance (Fig. 4b), resulting in a 13.4% (95% CI: 6.8 - 20.2%) reduction in maximal growth rate. This temperature dependent cost of resistance matched the effect of phage on bacteria growth (Fig. 3a and Fig. 5b). There was no difference in the optimum temperature between resistant and susceptible clones. Taken together, these results explain how ecological and evolutionary impacts of phage can alter the thermal performance of their host.

## Discussion

Here, we show that the presence of a parasite can profoundly impact the thermal performance of its host. Notably, phage reduced bacterial growth most at temperatures where the bacteria grew fastest, close to the bacterial *r*_*max*_, while having little or no impact at cold or high temperatures beyond *T*_*opt*_ (Fig. 1 & Fig. 2). This resulted in a higher optimum temperature for bacterial growth in the presence of phage (Fig. 2b). These results can be explained by a combination of ecological and evolutionary processes. At temperatures below the critical thermal maxima of the phage, phage epidemics vastly reduced bacterial abundance. In contrast, phage could not infect above 30 °C, but bacteria still had high growth rates, which resulted in a higher optimum temperature for the host in the presence of phage. However, rapid evolution also played an important role in altering the thermal performance of *P. fluorescens*. While phage resistance evolved rapidly at all temperatures below the phage *CT*_*max*_, at higher temperatures there was no selection for resistance (Fig. 3). Critically, there were costs associated with resistance, but these costs changed non-linearly with temperature (Fig. 4). At low temperatures and temperatures far beyond the bacteria *T*_*opt*_ there was no measurable cost of resistance, but significant costs of resistance at temperatures where bacteria growth was highest (Fig. 4). It is worth noting that costs of phage resistance were also greatest at the optimum temperature in another well studied bacteria-phage system; *Escherichia coli* and bacteriophage T4^25,26^.

How general are these results likely to be? We suggest that parasites (and symbionts more generally) impacts on host TPCs are likely widespread, because no change in host TPC would occur only when host and parasite traits respond equivalently with temperature. In reality, there are almost certainly mismatches between host and parasite TPCs, and rapid (co)evolutionary interactions between hosts and parasites, and differences in local adaptation to prevailing temperatures, appears to be the norm^6,27^. However, as with the effect of changing temperature on disease severity, precisely how TPCs will change will be context dependent. For example, we found the phage had a lower *CT*_*max*_ than the bacteria; a finding in contrast to recent work arguing that viruses and smaller organisms typically have broader thermal ranges and higher thermotolerances than their hosts^9,10^.

In conclusion, our study demonstrated that host-parasite interactions change in non-linear ways with temperature (G x G x E interaction), and this had a significant impact on the thermal performance of the host. By measuring the thermal performance of the host and the parasite simultaneously, and also examining the evolution and cost of resistance, we identified the mechanisms through which phage altered the thermal performance of the host. Our results highlight that TPCs measured under axenic conditions should be interpreted with caution; measuring TPCs in the absence of their parasites (and other associated microbiota) may not be reflective of the host’s TPC in nature where such interactions are ubiquitous. Future work should investigate the longer term evolutionary and coevolutionary consequences of climate warming^28^, and in a broader, more realistic ecological context, to determine how this impacts bacteria-phage interactions. In an era of human-induced climate change, it is more important than ever to gain a deeper understanding of how evolutionary and ecological processes can indirectly impact thermal performance and how host-parasite interactions will change with temperature.

## Materials and Methods

### Measuring bacterial growth in the presence and absence of phage

Isogenic *Pseudomonas fluorescens* SBW25 was cultured overnight (from a frozen stock) at 28 °C in 6 mL of M9 minimal salts media (M9), supplemented with 5 g of glycerol and 10 g of peptone (50 % concentration of King’s medium B) in glass vials at 180 r.p.m. Overnight stocks were then diluted to ∼ 50,000 cells per 10 µL (5 x 10^6^ cells per mL). Growth curves were measured in 96 well plates, with 180 µL of altered M9 (described above). We inoculated wells with 10 µL of bacteria and either 10 µL of M9 or 10 µL of phage (∼50 phage) giving a multiplicity of infection (MOI) of 0.001. We left 6 wells free for both bacteria and bacteria plus phage treatments at each temperature as blank controls. We set up 6 replicates of bacteria and bacteria plus phage simultaneously at 8 temperatures (15, 20, 25, 28, 30, 33, 35 and 37 °C). Each plate was placed in a plastic box with a moist sponge at the bottom to prevent evaporation of media from the wells which may confound measurements of optical density (OD). OD (600 nm wavelength) was measured as a proxy for density of *Pseudomonas fluorescens* using a plate reader (Biotek Instruments Ltd). Readings of OD were taken with the lid off at an average of every 3 hours for up to 75 hours.

### Measuring phage replication rate

Replication of the phage SBWΦ2 was measured, using methods similar to Knies *et al.*^29,30^, at the same temperatures as the bacterial growth curves, with the addition of 3 additional temperatures (22.5 °C, 26 °C and 27 °C) to better characterise temperatures around the optimum of phage replication. First, isogenic *P. fluorescens* was grown overnight in conditions described above. The bacteria were transferred into fresh media at 28 °C and allowed to grow for 6 hours while shaking to increase density. We then added 20 µL of phage (∼ 10^6^) to each tube (six replicates per temperature). Vials were left static for 4 hours at each temperature, after which phage was extracted using chloroform extraction. 100 µL of chloroform was added to 900 µL of culture, then vortexed and centrifuged at 10000 g for 5 minutes. The supernatant was removed and placed in fresh Ependorf tubes. Phage titres were measured using plaque assays against the ancestral bacteria at 28 °C.

### Measuring resistance of bacteria

To investigate the mechanism behind any effect of phage on bacterial growth, we measured the resistance of bacteria within a single growth curve. We set up 18 wells of 96 well plates at 8 temperatures that contained ∼50,000 cells and ∼50 phage (as described above). We then destructively sampled 6 wells at three time points through the growth curve (after 12, 24 and 48 hours). To do this, 20 µL of each well was placed in 180 µL of M9. These were then serially diluted and plated onto KB agar. Twelve colonies from each replicate were taken per time point and grown overnight in 150 µL of altered M9, shaking at 28 °C. Each clone was then checked for resistance against the ancestral phage using a phage streak assay. Phage streak assays were incubated overnight at 28 °C.

### Measuring the cost of resistance

To determine whether any effect of phage was due to a cost of resistance, we grew 12 replicates of *P. fluorescens* in the presence and absence of phage for 12 hours at 28 °C. After 12 hours, each population was plated onto KB agar and grown for 2 days at 28 °C. Three clones were isolated from each replicate and grown for two days in modified M9 media. Each isolate was checked for resistance against the ancestral phage. Growth curves of each clone were done using the methods described above, but inoculate density was ∼500,000 cells to reduce the lag phase and no phage was added.

## Statistical analyses

### Calculating exponential growth rate for bacteria

For bacterial growth, we wanted to estimate exponential population growth rate in the presence and absence of phage, and for resistant and susceptible clones. In the presence and absence of phage, prior to model fitting we manually removed points which deviated from the expected shape of logistical growth. These anomalies were likely caused by decreases in abundance due to phage infection. When these fluctuations occurred in the exponential growth phase, they led to exponential growth rate being underestimated, preventing the comparison between the exponential growth rate of bacteria in the presence of phage at some temperatures. After this initial data cleaning, we fitted the Gompertz model^31^ to measurements of *log*_*10*_*OD*_*600*_ through time, *t*, in hours, using code extracted from the *R* package ‘*nlsMicrobio*’^32^:

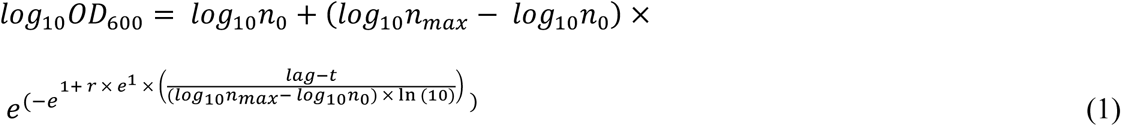

Where *log*_*10*_*n*_*0*_ is the starting density, *log*_*10*_*n*_*max*_ is carrying capacity, *r* is the exponential growth rate (hr^-1^) and *lag* is the lag time in hours. Model fitting was done using nonlinear least squares regression using the *R* package ‘*nls.multstart*’^33^. This method of model fitting involved running up to 500 iterations of the fitting process with start parameters drawn from a uniform distribution and retaining the fit with the lowest Akaike Information Criterion score (AIC). The parameters of the model (*r, log*_*10*_*n*_*0*_, *log*_*10*_*n*_*max*_ and *lag*) can be seen as growth ‘traits’ which may vary with both temperature and the presence and absence of phage. Other growth models were fitted (e.g. Baranyi, Buchanan; Table S3), but the Gompertz model returned lower AIC scores for the majority of model fits (Figure S1 & Figure S2).

For susceptible and resistant clones, we cleaned the data by removing the first time point where bubbles were likely and setting time zero to the time at which the first optical density measurement was detected for each clone. We then initially used the same modelling approach, but this time the Baranyi model without lag was the model most selected using AIC scores (Figure S4). However, after examining the predictions and residuals of the model fits (Figure S5, Figure S6), we found that exponential growth rate was underestimated at temperatures where bacteria grew best, and at these temperatures there was a significantly greater underestimation of growth rate in susceptible, rather than resistant, bacteria (Figure S7). Consequently, exponential growth rate per clone was calculated here using rolling regression, taking the steepest slope of the linear regression between *lnOD*_*600*_ and time in hours in a shifting window of every 4 time points as the estimate of exponential growth. Average growth rate per replicate was calculated by taking the mean clonal growth rate. After data cleaning and model fitting, every growth curve had estimates of exponential growth rate which were then used to model the thermal performance of bacteria.

### Fitting thermal performance curves to phage and bacteria

Thermal performance curves were fitted for phage replication rate and *r* of bacteria in the presence and absence of phage, and for resistant and susceptible bacterial clones. We used the Sharpe-Schoolfield equation for high-temperature inactivation^34^, which extends the original Boltzmann equation to incorporate a decline in growth rate beyond the optimum.

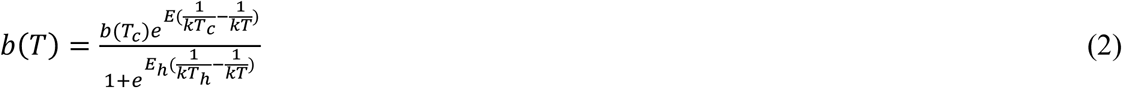

*b*(*T*) is the rate of phage replication or bacterial growth at temperature, *T*, in Kelvin (K). Instead of the intercept being at 0 K (-273.15 °C), *b*(*T*_c_) is the rate at a common temperature, *T*_c_ = 20 °C (293.15 K)^35^. *E* (eV) is the activation energy that describes the temperature-dependence of the biological rate, *k* is Boltzmann’s constant (8.62 × 10^-5^ eV K^-1^), *E*_*h*_ (eV) characterises the high temperature-induced inactivation of enzyme kinetics and *T*_*h*_ (K) is the temperature at which half the rate is inactivated due to high temperatures. Equation 2 yields an optimum temperature, *T*_*opt*_, (K).

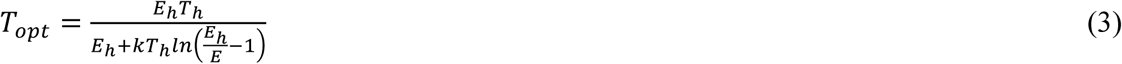

Maximal growth rate, *r*_*max*_, was calculated by using the estimated model parameters to predict the rate at *T*_*opt*_. As in previous studies^16,17^, these ‘traits’ were then used to look for differences across treatments. As phage replication was negative at high temperatures, an offset was added to the equation to raise all rates above 0 to allow model fitting. This invalidated any interpretation of the activation energy and deactivation energy for phage replication, but this is difficult already. However, this was already difficult as phage replication is partially determined by bacteria growth rate which is also temperature dependent. Consequently, for phage replication we concentrated on the optimum temperature (*T*_*opt*_) and critical thermal maximum (*CT*_*max*_) which is the temperature at which phage replication became negative at high temperatures.

For phage and bacteria, Equation 2 was fitted to the data using non-linear regression in a Bayesian framework using the *R* package ‘*brms*’^36^. This method allows for prior information on suitable parameter values and the estimation of uncertainty around predictions and parameters, including derived parameters not present in the original model formulation such as *T*_*opt*_, *CT*_*max*_ and *r*_*max*_. Different models were fitted for phage replication rate, exponential growth rate of bacteria in the presence and absence of phage, and exponential growth rate of resistant and susceptible bacterial clones. For the analysis including resistant and susceptible clones, a random effect was added to account for the non-independence of measurements of the same population across temperatures. For bacteria exponential growth rate, phage presence/absence or susceptible/resistance was added as a factor that could alter each parameter of the model. Models were run for 5000 iterations and 3 chains were used with uninformative priors. Model convergence was assessed using posterior predictive checks, Rhat values (all values were 1) and manually checking of chain-mixing. Differences between parameter estimates are described using 95% credible intervals. Credible intervals of predictions and parameters were calculated from the posterior distribution using the *R* package ‘*tidybayes*’^37^. Non-overlapping 95% credible intervals indicate statistical significance at (at least) the p = 0.05 level.

Using predictions from the model for bacterial growth, the relative fitness of bacteria in the presence of phage was estimated across the continuous temperature range (15 – 37 °C). The difference was calculated as a selection coefficient, where relative fitness at each temperature, *w*(*T*), was calculated as:

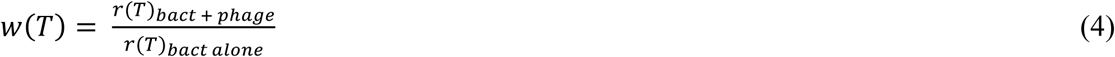

where *r*(*T*)_*bact+phage*_ is the growth rate at a given temperature in the presence of phage and *r*(*T*)_*bact alone*_ is the growth rate in the absence of phage. When the 95% credible intervals of the predictions do not cross 1, it indicates that phage significantly altered bacterial growth rate. When there is overlap with the predictions and 1, it means there is no significant change in relative fitness. An identical statistical approach was taken for analysing the growth rates of susceptible and resistant clones. In this instance, the relative fitness across temperatures, *w*(*T*), represented the cost of resistance.

### Analysing phage resistance assays

A logistic regression was used to analyse the proportion of resistance through time and across temperatures. A binomial model was fitted to the number of resistant and susceptible individuals per replicate at each temperature and time point using the logit transformation. As there were many populations where all clones were completely susceptible or resistant (resulting in zero and one inflated data), we added one to both the number of resistant and susceptible individuals in each population and used a quasibinomial error structure to control for overdispersion. This data transformation reduced the variance in estimates which resulted in conservative estimates of differences in resistance between temperatures and through time. We fitted a model that combined the number of resistant and susceptible clones in a population as the response variable and included temperature and time (in hours) as discrete predictor variables. Model selection was done through likelihood ratio tests using *F* tests. Pairwise post-hoc comparisons were done on the response scale using the *R* package ‘*emmeans’*^38^. All analyses were done using the statistical programming language *R* (v3.5.1)^39^ and all plots were made using the *R* package ‘*ggplot2*’^40^.

## Acknowledgements

This work was funded by a NERC grant awarded to AB.

## References

1. Harvell, C. D. et al. Climate warming and disease risks for terrestrial and marine biota. Science 296, 2158–2162 (2002).

2. Demory, D. et al. Temperature is a key factor in Micromonas–virus interactions. The ISME journal 11, 601 (2017).

3. Paull, S. H., LaFonte, B. E. & Johnson, P. T. Temperature-driven shifts in a host-parasite interaction drive nonlinear changes in disease risk. Global Change Biology 18, 3558–3567 (2012).

4. Paull, S. H., Raffel, T. R., LaFonte, B. E. & Johnson, P. T. How temperature shifts affect parasite production: testing the roles of thermal stress and acclimation. Functional Ecology 29, 941–950 (2015).

5. Molnár, P. K., Kutz, S. J., Hoar, B. M. & Dobson, A. P. Metabolic approaches to understanding climate change impacts on seasonal host-macroparasite dynamics. Ecology Letters 16, 9–21 (2013).

6. Gehman, A.-L. M., Hall, R. J. & Byers, J. E. Host and parasite thermal ecology jointly determine the effect of climate warming on epidemic dynamics. Proceedings of the National Academy of Sciences 201705067 (2018).

7. Cohen, J. M. et al. The thermal mismatch hypothesis explains host susceptibility to an emerging infectious disease. Ecology letters 20, 184–193 (2017).

8. Nowakowski, A. J. et al. Infection risk decreases with increasing mismatch in host and pathogen environmental tolerances. Ecology letters 19, 1051–1061 (2016).

9. Mojica, K. D. & Brussaard, C. P. Factors affecting virus dynamics and microbial host–virus interactions in marine environments. FEMS microbiology ecology 89, 495–515 (2014).

10. Rohr, J. R. et al. The complex drivers of thermal acclimation and breadth in ectotherms. Ecology letters 21, 1425–1439 (2018).

11. Thomas, M. B. & Blanford, S. Thermal biology in insect-parasite interactions. Trends in Ecology & Evolution 18, 344–350 (2003).

12. Sinclair, B. J. et al. Can we predict ectotherm responses to climate change using thermal performance curves and body temperatures? Ecology Letters 19, 1372–1385 (2016).

13. Sunday, J. M., Bates, A. E. & Dulvy, N. K. Thermal tolerance and the global redistribution of animals. Nature Climate Change 2, 686–690 (2012).

14. Luhring, T. M. & DeLong, J. P. Predation changes the shape of thermal performance curves for population growth rate. Current zoology 62, 501–505 (2016).

15. Tseng, M. & O’Connor, M. I. Predators modify the evolutionary response of prey to temperature change. Biology letters 11, 20150798 (2015).

16. Padfield, D., Yvon-Durocher, G., Buckling, A., Jennings, S. & Yvon-Durocher, G. Rapid evolution of metabolic traits explains thermal adaptation in phytoplankton. Ecology Letters 19, 133–142 (2016).

17. Schaum, C.-E. et al. Adaptation of phytoplankton to a decade of experimental warming linked to increased photosynthesis. Nature Ecology & Evolution 1, 0094 (2017).

18. Angilletta Jr, M. J. Thermal adaptation: a theoretical and empirical synthesis. (Oxford University Press, 2009).

19. Lopez-Pascua, L.D.C & Buckling, A. Increasing productivity accelerates host–parasite coevolution. Journal of evolutionary biology 21, 853–860 (2008).

20. Buckling, A. & Rainey, P. B. Antagonistic coevolution between a bacterium and a bacteriophage. Proceedings of the Royal Society of London B: Biological Sciences 269, 931–936 (2002).

21. Gómez, P. & Buckling, A. Bacteria-phage antagonistic coevolution in soil. Science 332, 106–109 (2011).

22. Zhang, Q.-G. & Buckling, A. Antagonistic coevolution limits population persistence of a virus in a thermally deteriorating environment. Ecology letters 14, 282–288 (2011).

23. Buckling, A. & Rainey, P. B. The role of parasites in sympatric and allopatric host diversification. Nature 420, 496 (2002).

24. Lenski, R. E. & Levin, B. R. Constraints on the coevolution of bacteria and virulent phage: a model, some experiments, and predictions for natural communities. The American Naturalist 125, 585–602 (1985).

25. Quance, M. A. & Travisano, M. Effects of temperature on the fitness cost of resistance to bacteriophage T4 in Escherichia coli. Evolution: International Journal of Organic Evolution 63, 1406–1416 (2009).

26. Cooper, V. S., Bennett, A. F. & Lenski, R. E. Evolution of thermal dependence of growth rate of Escherichia coli populations during 20,000 generations in a constant environment. Evolution 55, 889–896 (2001).

27. Dell, A. I., Pawar, S. & Savage, V. M. Systematic variation in the temperature dependence of physiological and ecological traits. Proceedings of the National Academy of Sciences 108, 10591–10596 (2011).

28. Gorter, F. A., Scanlan, P. D. & Buckling, A. Adaptation to abiotic conditions drives local adaptation in bacteria and viruses coevolving in heterogeneous environments. Biology letters 12, 20150879 (2016).

29. Knies, J. L., Izem, R., Supler, K. L., Kingsolver, J. G. & Burch, C. L. The genetic basis of thermal reaction norm evolution in lab and natural phage populations. PLoS biology 4, e201 (2006).

30. Knies, J. L., Kingsolver, J. G. & Burch, C. L. Hotter is better and broader: thermal sensitivity of fitness in a population of bacteriophages. The American Naturalist 173, 419–430 (2009).

31. Gompertz, B. On the nature of the function expressive of the law of human mortality, and on a new mode of determining the value of life contingencies. Philosophical Transactions of the Royal Society of London B: Biological Sciences 115, 513–583 (1825).

32. Baty, F. & Delignette-Muller, M. L. nlsMicrobio: Nonlinear regression in predictive microbiology. R package version 3.2. 2. (2016).

33. Padfield, D. & Matheson, G. nls.multstart: Robust Non-Linear Regression using AIC Scores. R package version 1.0.0. (2018).

34. Schoolfield, R. M., Sharpe, P. J. H. & Magnuson, C. E. Non-linear regression of biological temperature-dependent rate models based on absolute reaction-rate theory. Journal of theoretical biology 88, 719–731 (1981).

35. Padfield, D., Buckling, A., Warfield, R., Lowe, C. & Yvon-Durocher, G. Linking phytoplankton community metabolism to the individual size distribution. Ecology Letters (2018).

36. Bürkner, P.-C. brms: An R package for Bayesian multilevel models using Stan. Journal of Statistical Software 80, 1–28 (2017).

37. Kay, M. tidybayes: Tidy Data and Geoms for Bayesian Models. (2018).

38. Lenth, R. Emmeans: Estimated marginal means, aka least-squares means. R Package Version 1, (2018).

39. Team, R. C. R: A language and environment for statistical computing. (2013).

40. Wickham, H. ggplot2: elegant graphics for data analysis. (Springer, 2016).

