## Supplementary Information for "Temperature-dependent changes to host-parasite interactions alter the thermal performance of a bacterial host"

**Author contributions:** D.P and A.B conceived the study and designed the experimental work. D.P and M.C conducted the experiments. D.P analysed the data. All authors contributed significantly to the first draft of the manuscript and to revisions.

**Data accessibility statement:** All data and R code used in the analysis will be made available on GitHub and archived on Zenodo.

**Table S1. Point estimates and 95% credible intervals (as determined using Bayesian methods) for fitted and derived metabolic traits.**

| Rate | Parameter | Mean | 2.5% | 97.5% |
|---|---|---|---|---|
| phage replication | $CT_{max}$ (ºC) | 29.2 | 29 | 29.4 |
| | $T_{opt}$ (ºC) | 27.0 | 26.5 | 27.5 |
| bacteria growth without phage | E (eV) | 0.84 | 0.59 | 1.16 |
| | $E_h$ (eV) | 2.36 | 2.03 | 2.79 |
| | $T_{opt}$ (ºC) | 28.0 | 27.1 | 29.0 |
| | $r_{max}$ ($hr^{-1}$) | 0.72 | 0.68 | 0.76 |
| bacteria growth with phage | E (eV) | 0.33 | 0.20 | 0.50 |
| | $E_h$ (eV) | 4.25 | 2.57 | 6.63 |
| | $T_{opt}$ (ºC) | 30.6 | 29.0 | 32.1 |
| | $r_{max}$ ($hr^{-1}$) | 0.57 | 0.54 | 0.62 |
| bacteria growth of susceptible clones | E (eV) | 0.49 | 0.42 | 0.57 |
| | $E_h$ (eV) | 2.32 | 1.95 | 2.77 |
| | $T_{opt}$ (ºC) | 30.5 | 30.0 | 31.0 |
| | $r_{max}$ ($hr^{-1}$) | 0.77 | 0.73 | 0.81 |
| bacteria growth of resistant clones | E (eV) | 0.42 | 0.33 | 0.56 |
| | $E_h$ (eV) | 1.95 | 1.47 | 2.57 |
| | $T_{opt}$ (ºC) | 30.2 | 29.2 | 31.0 |
| | $r_{max}$ ($hr^{-1}$) | 0.66 | 0.63 | 0.70 |
| bacteria growth | % change in $r_{max}$ due to presence of phage | -20.6 | -13.1 | -27.3 |
| | % change in $r_{max}$ due to phage resistance | -13.6 | -6.8 | -20.2 |

Parameters include $CT_{max,}$ the critical thermal maximum, $T_{opt}$, the optimum temperature, E, the activation energy, $E_h$, the deactivation energy, $r_{max}$, the maximum growth rate and the % change in maximum growth rate due to phage presence and due to phage resistance. Not all parameters are shown for each rate because they were either outside the range of the data collected or were not biologically meaningful for the data collected.

**Table S2. Results of multiple pairwise comparisons between resistance through time at each temperature.**

| Temperature | Contrast | Odds ratio | SE | z ratio | p value |
|---|---|---|---|---|---|
| 15 | 12 hours vs. 24 hours | 1 | 0.43 | 0 | 1 |
| | 12 hours vs. 48 hours | 1.26 | 0.54 | 0.54 | 0.85 |
| | 24 hours vs. 48 hours | 1.26 | 0.54 | 0.54 | 0.85 |
| 20 | 12 hours vs. 24 hours | 1.04 | 0.45 | 0.08 | 0.99 |
| | 12 hours vs. 48 hours | 1.01 | 0.44 | 0.03 | 0.99 |
| | 24 hours vs. 48 hours | 0.98 | 0.43 | -0.05 | 0.99 |
| 25 | 12 hours vs. 24 hours | 1.16 | 0.62 | 0.89 | 0.65 |
| | 12 hours vs. 48 hours | 5.61 | 2.05 | 4.71 | <0.001 |
| | 24 hours vs. 48 hours | 3.85 | 1.29 | 4.02 | <0.001 |
| 28 | 12 hours vs. 24 hours | 1.62 | 0.66 | 1.17 | 0.47 |
| | 12 hours vs. 48 hours | 13.9 | 4.96 | 7.36 | <0.001 |
| | 24 hours vs. 48 hours | 8.60 | 2.81 | 6.59 | <0.001 |
| 30 | 12 hours vs. 24 hours | 0.02 | 0.01 | -10.46 | <0.001 |
| | 12 hours vs. 48 hours | 0.37 | 0.16 | -2.35 | 0.049 |
| | 24 hours vs. 48 hours | 16.8 | 5.48 | 8.68 | <0.001 |
| 33 | 12 hours vs. 24 hours | 0.92 | 0.41 | -0.19 | 0.98 |
| | 12 hours vs. 48 hours | 1 | 0.43 | 0 | 1 |
| | 24 hours vs. 48 hours | 1.1 | 0.49 | 0.194 | 0.98 |
| 35 | 12 hours vs. 24 hours | 0.85 | 0.35 | -0.40 | 0.92 |
| | 12 hours vs. 48 hours | 0.99 | 0.43 | -0.03 | 0.99 |
| | 24 hours vs. 48 hours | 1.12 | 0.49 | 0.37 | 0.93 |
| 37 | 12 hours vs. 24 hours | 1 | 0.433 | 0 | 1 |
| | 12 hours vs. 48 hours | - | - | - | - |
| | 24 hours vs. 48 hours | - | - | - | - |

At temperatures where growth was highest, resistance changed significantly through time. P values were adjusted using the tukey method for comparing a family of 3 estimates and tests were performed on the log odds ratio scale. An odds ratio of 1 would indicate that resistance was the same in both groups, with a higher odds ratio indicating that resistance was higher in the first group, and a lower odds ratio would indicate that resistance was higher in the second group.

**Table S3. Logistical growth equations used in the modelling of bacterial growth in the presence and absence of phage.**

| Model | Equation |
|---|---|
| Gompertz | $log_{10}OD_{600} = log_{10}n_0 + (log_{10}n_{max} - log_{10}n_0) \times e^{(-e^{1+r \times e^1 \times \left(\frac{lag-t}{(log_{10}n_{max} - log_{10}n_0) \times \ln(10)}\right)})}$ |
| Baranyi | $log_{10}OD_{600} = log_{10}n_{max} + log_{10}(\frac{-1 + e^{r \times lag} + e^{r \times t}}{e^{r \times t} - 1 + e^{r \times lag} \times 10^{(log_{10}n_{max} - log_{10}n_0)}})$ |
| Baranyi without lag | $log_{10}OD_{600} = log_{10}n_{max} - log_{10}\left(1 + (10^{(log_{10}n_{max} - log_{10}n_0)} - 1\right) \times e^{-r \times t})$ |
| Buchanan | $log_{10}OD_{600} = log_{10}n_0$ for when $t \leq lag$<br>$log_{10}OD_{600} = log_{10}n_0 + r(t - lag)$ for when $lag \leq t \leq t_s$<br>$log_{10}OD_{600} = log_{10}n_{max}$ for when $t \geq t_s$ |
| Buchanan without lag | $log_{10}OD_{600} = log_{10}n_0 + r(t - lag)$ for when $t \leq t_s$<br>$log_{10}OD_{600} = log_{10}n_{max}$ for when $t \geq t_s$ |

Where $log_{10}OD_{600}$ is the log10 of the absorbance measurement, $log_{10}n_0$ is the starting density, $log_{10}n_{max}$ is carrying capacity, $r$ is the exponential growth rate ($hr^{-1}$), $lag$ is the lag time in hours and $t_s$ is the time to stationary phase in hours. Model equations were copied from the R package '*nlsMicrobio*'. Code for fitting each equation and comparing AIC scores can be found on the GitHub repository for this manuscript.

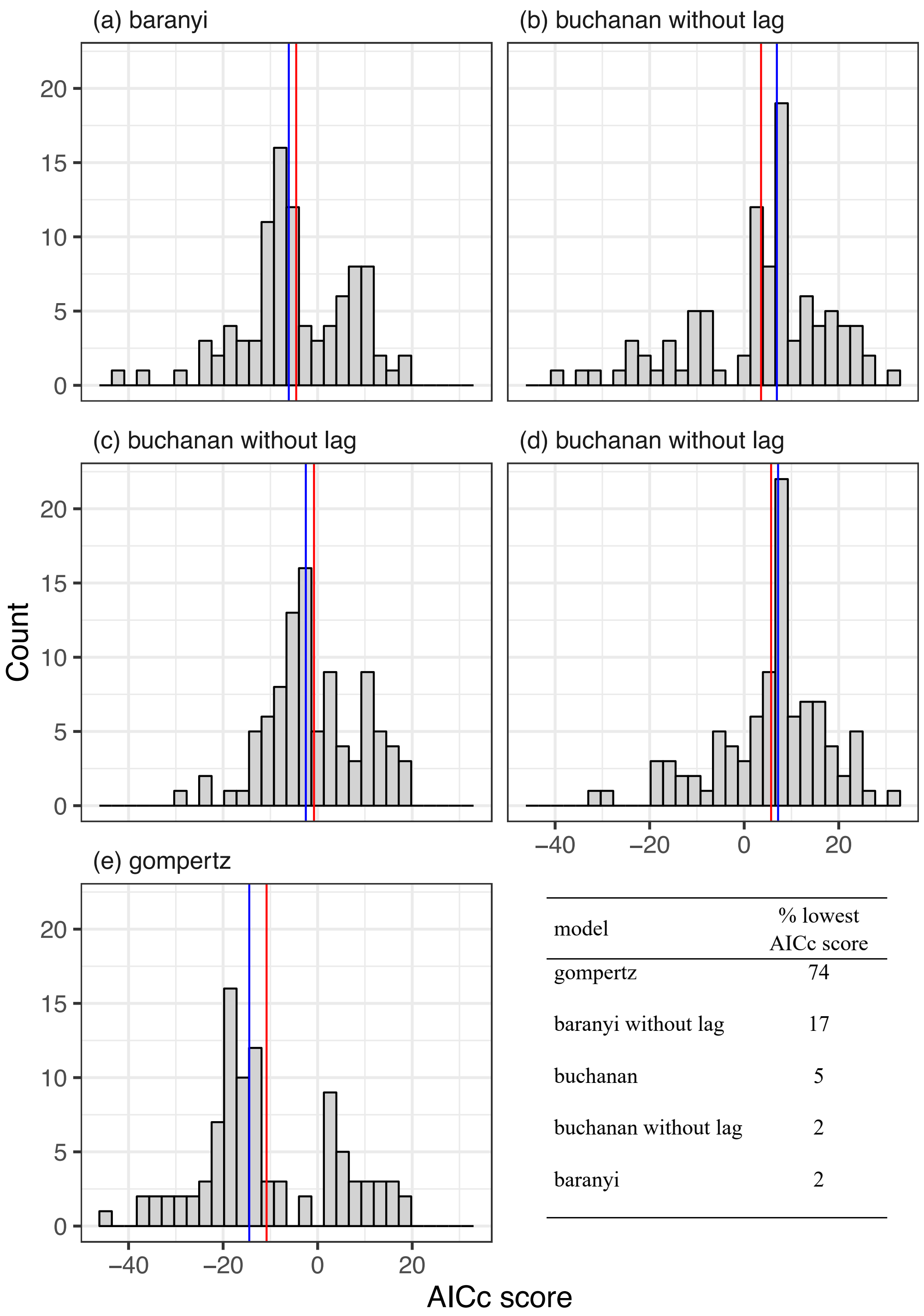


| model | % lowest AICc score |
| --- | --- |
| gompertz | 74 |
| baranyi without lag | 17 |
| buchanan | 5 |
| buchanan without lag | 2 |
| baranyi | 2 |

**Figure S1. Distribution of AICc scores for different logistical growth models fitted to bacteria growth in the presence and absence of phage.** Numerous logistical growth models were fitted to each bacterial growth curve in the presence and absence of phage. The Akaike's Information Criterion score adjusted for small samples (AICc) for each model was calculated and compared across models to select the best, consensus model. The table in the bottom right demonstrates that for 74% of the curves, the Gompertz model returned the lowest AICc score. The red and blue lines per panel represent the mean and median AICc score of that model respectively.

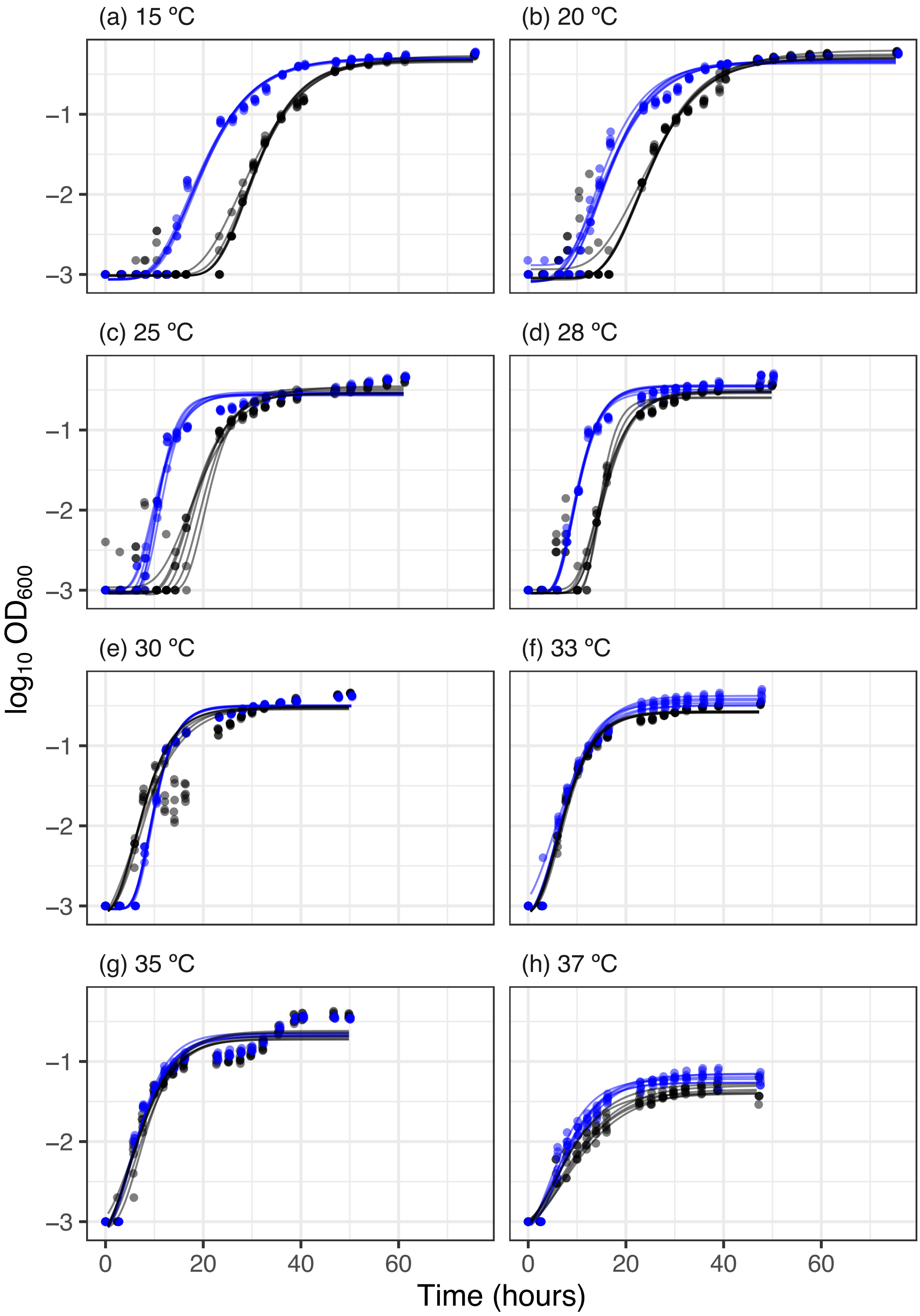


**Figure S2. Logistic growth curves for bacterial growth in the presence (black) and absence (blue) of phage.** The Gompertz model for logistic growth was fitted to each independent replicate and the exponential growth parameter was extracted for use in the thermal performance curves. Extreme abundance deviations can be seen in the presence of phage (black points) where phage infection is occurring. Points represent individual measurements and lines represent predictions of the best fitting model for each replicate at each temperature.

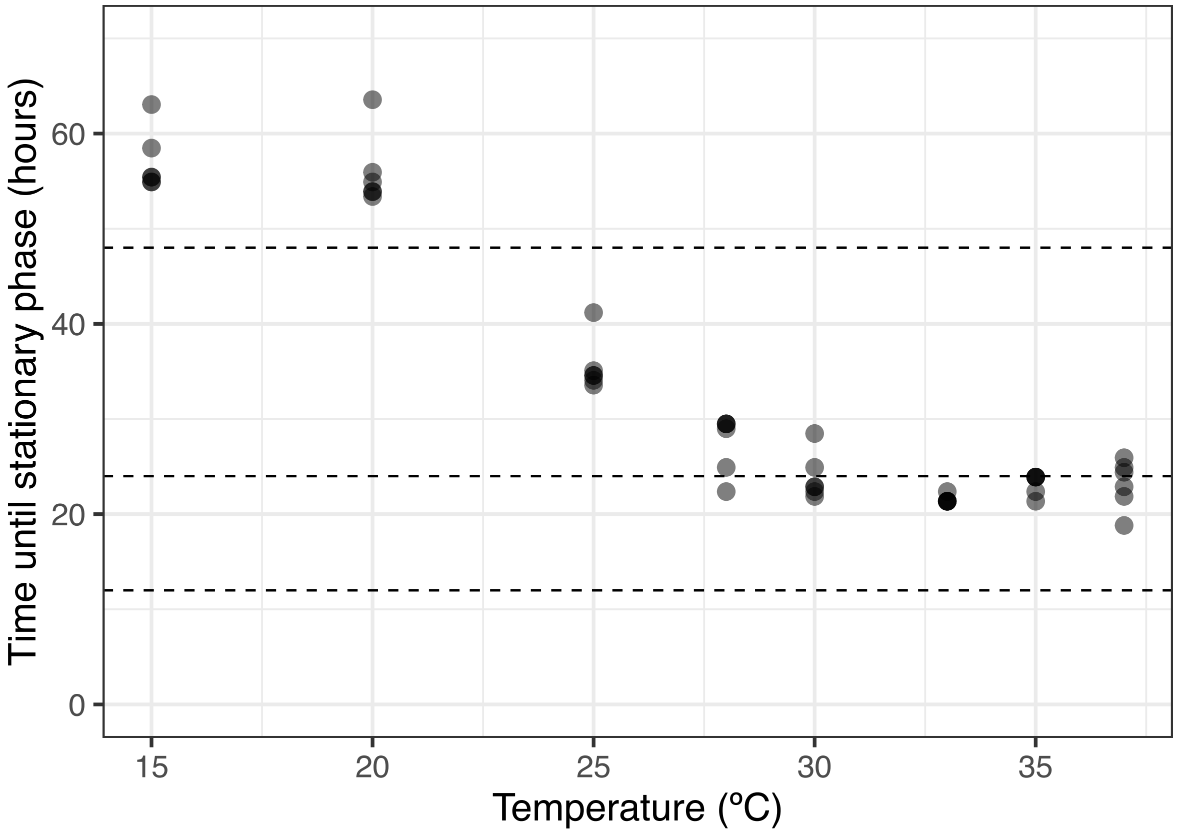


**Figure S3. Time to stationary phase of bacteria growth in the presence of phage across temperatures.** Time to stationary phase was estimated as the time at which the predictions of the model were 90% of the estimated carrying capacity, $log_{10}n_{max}$. Temperatures above 20 ºC are all in stationary phase before the final sampling point of 48 hours, indicating nutrient limitation between 24 and 48 hours at these temperatures. Points represent the time to stationary phase of independent replicates. Dashed lines indicate the times at which samples were taken to test for resistance in equivalent trials.

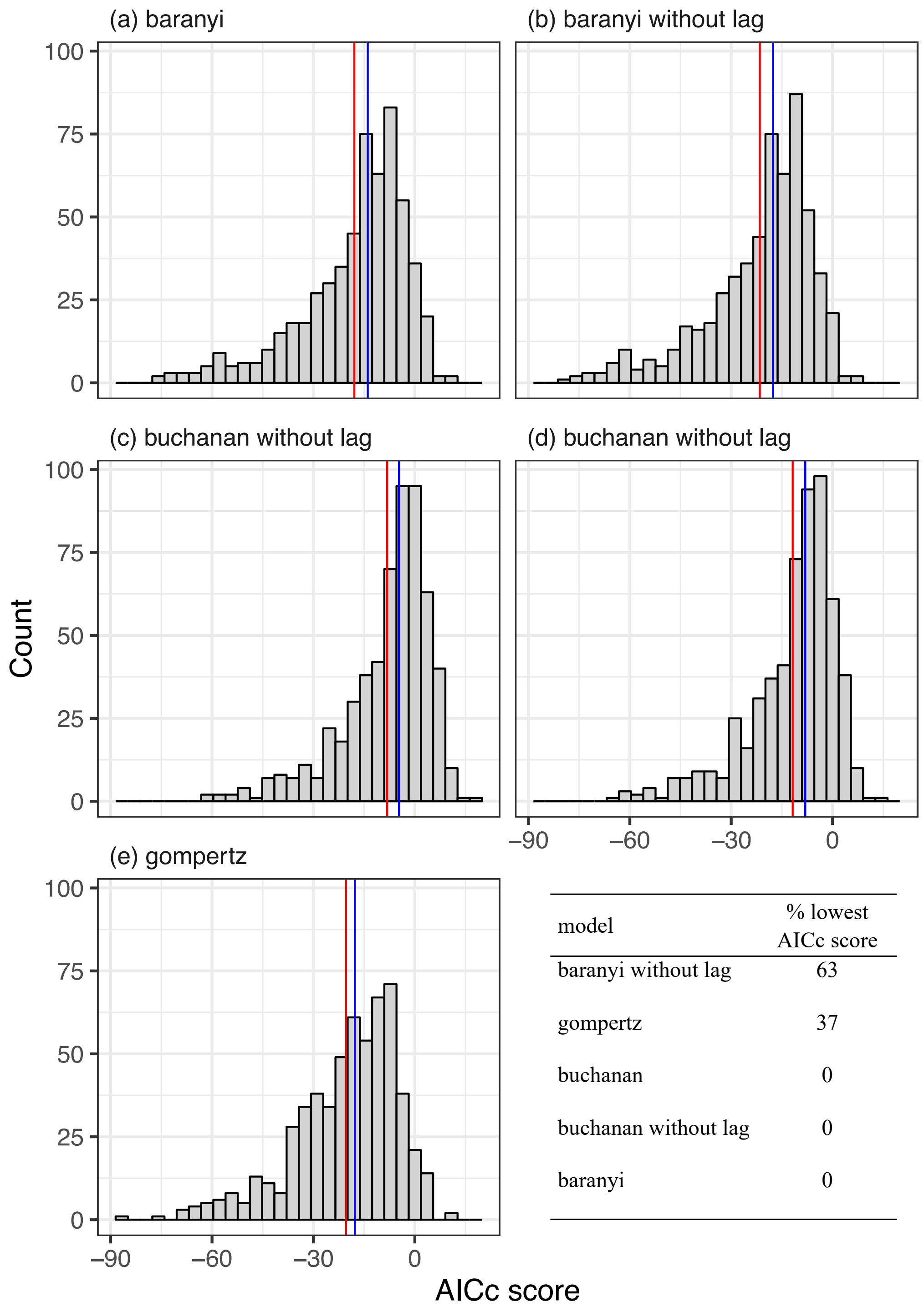


| model | % lowest AICc score |
|---|---|
| baranyi without lag | 63 |
| gompertz | 37 |
| buchanan | 0 |
| buchanan without lag | 0 |
| baranyi | 0 |

**Figure S4. Distribution of AICc scores for different logistical growth models fitted to bacteria growth of susceptible and resistant clones.** Numerous logistical growth models were fitted to each bacterial growth curve in the presence and absence of phage. The Akaike's Information Criterion score adjusted for small samples (AICc) for each model was calculated and compared across models to select the best, consensus model. The table in the bottom right demonstrates that for 63% of the curves, the Baranyi model without a lag phase returned the lowest AICc score. The red and blue lines per panel represent the mean and median AICc score of that model respectively.

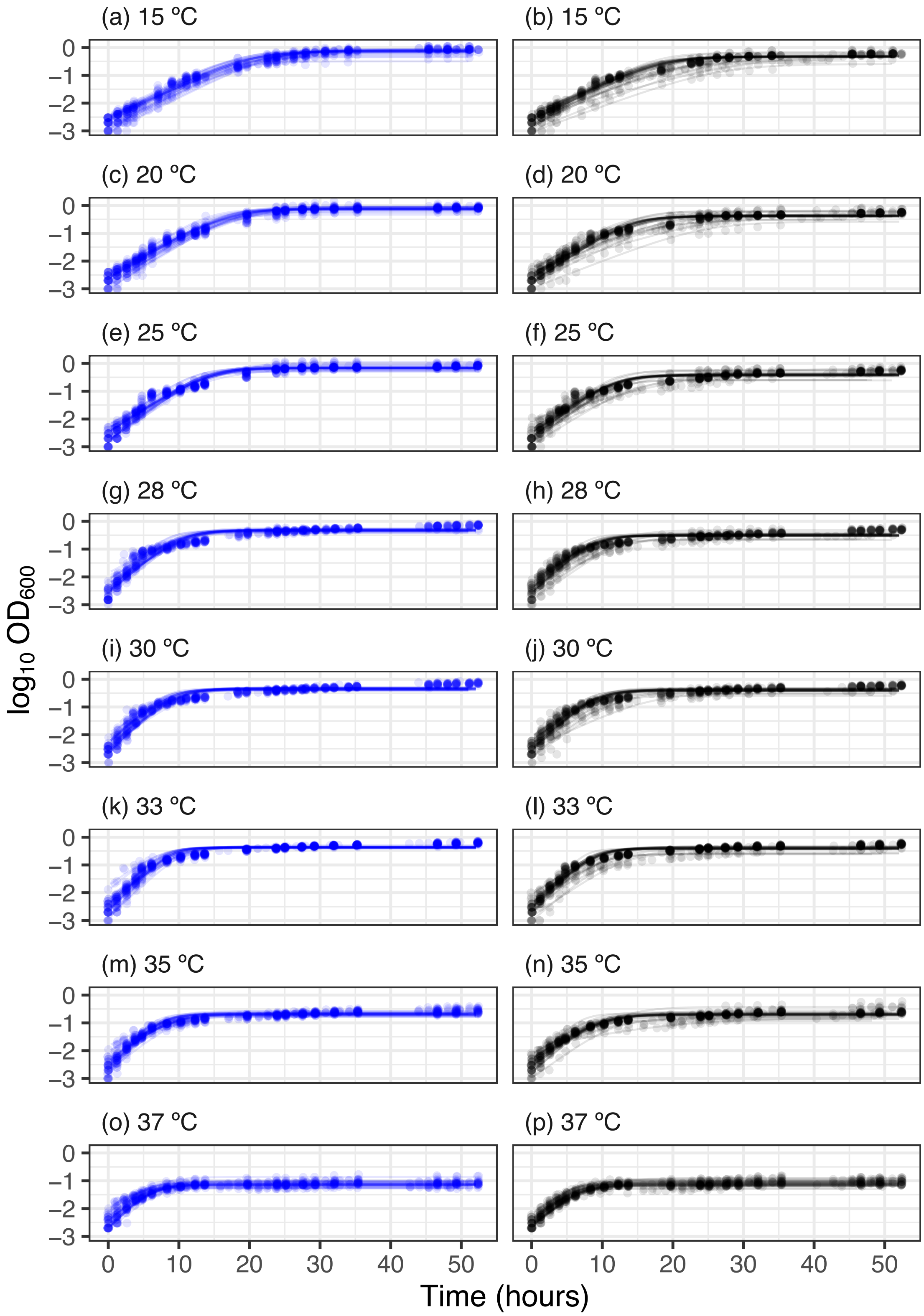


**Figure S5. Logistic growth curves for bacterial growth of susceptible (blue) and resistant (black) clones.** The Baranyi model without a lag phase was fitted to each independent replicate and the exponential growth parameter was extracted for use in the thermal performance curves. Extreme abundance deviations can be seen in the presence of phage (black points) where phage infection is occurring. Points represent individual measurements and lines represent predictions of the best fitting model for each replicate at each temperature.

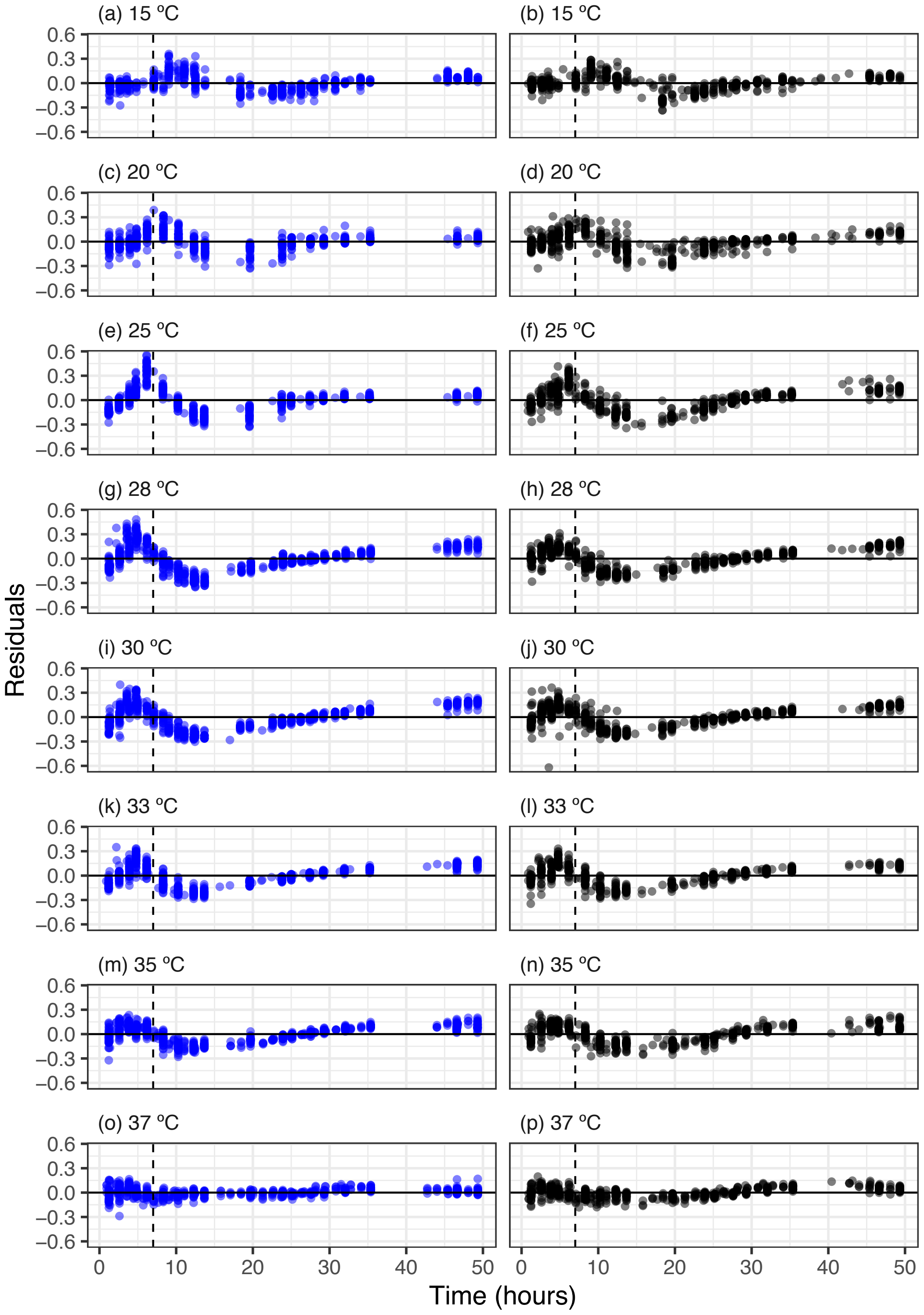


**Figure S6. Fit residuals through time of logistic growth curves of susceptible (blue) and resistant (black) bacterial clones.** The residuals of the Baranyi model without a lag phase were plotted as a function of time for each clone. There is some systematic variation in the residuals that are similar across most temperatures and resistant and susceptible clones. However, there does appear to be systematic variation in the first 7 hours after growth was first measured which could result in growth being underestimated at some temperatures more than others. The vertical line is drawn after 7 hours after growth was first detected and is a key portion of the curve used to estimate exponential growth rate.

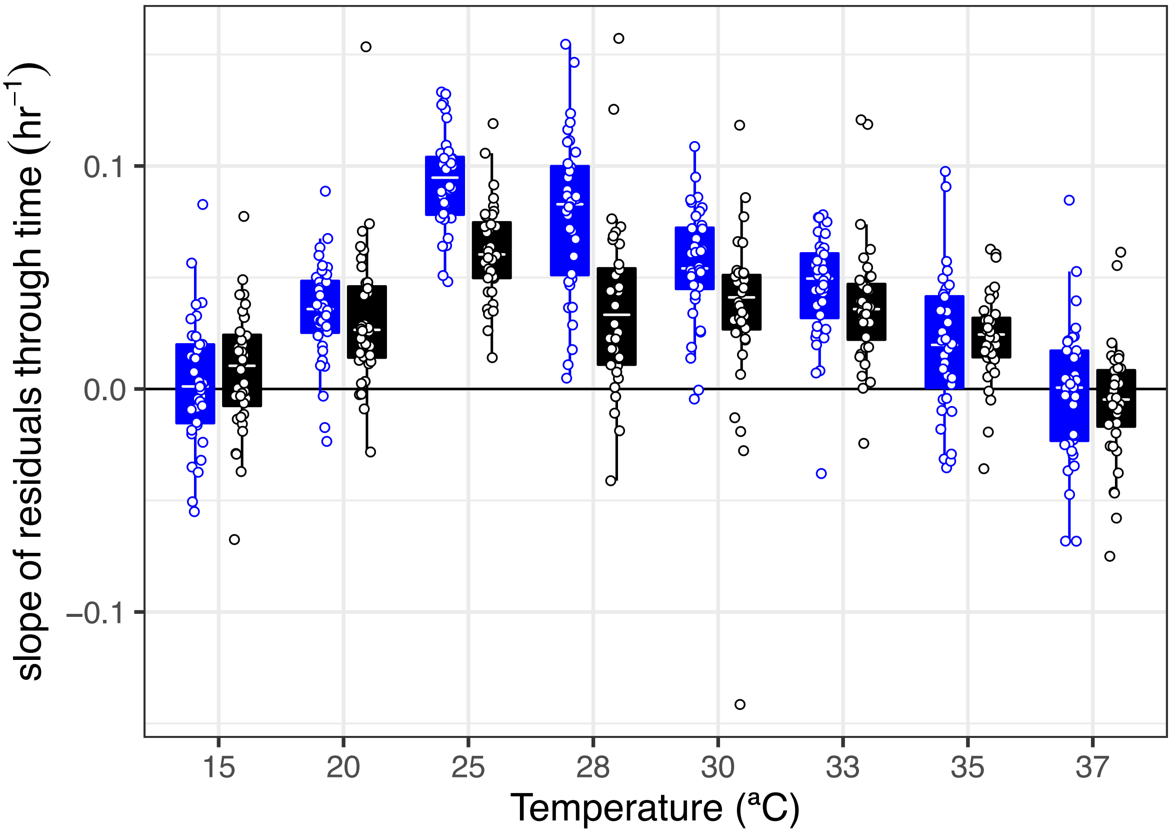


**Figure S7. Systematic variation in the residuals during the exponential growth phase of susceptible (blue) and resistant (black) bacterial clones.** The slope between the residuals and time over the first 7 hours growth was detected was investigated. A slope of 0 would indicate that the model estimates exponential growth adequately, whereas a slope greater than 1 would indicate that the model underestimates growth rate given the data. Exponential growth rate is underestimated at temperatures where bacteria grew best, and at these temperatures there was a significantly greater underestimation of growth rate in susceptible, rather than resistant bacteria. Points represent the slope of individual fits. Tops and bottoms of the bars represent the 75th and 25th percentiles of the data, the white lines are the medians, and the whiskers extend from their respective hinge to the smallest or largest value no further than 1.5 * interquartile range.
